# *Enterobacter Sp. SM3* Exhibits Run-and-Tumble Motility

**DOI:** 10.1101/2023.07.24.550425

**Authors:** Silverio Johnson, Brian Freedman, Jay X. Tang

## Abstract

The recent discovery of the peritrichous, swarm-competent bacterium *Enterobacter sp. SM3* has offered a new opportunity to elucidate the role of swarming motility in the gut microbiome. Here we present first findings of the run-and-tumble behavior of *SM3* in both a control solution of motility buffer and upon bulk exposure to the chemoattractants serine and aspartate, drawing a comparison with the well-studied behavior of *E. coli*. It was found that *SM3* runs with an average speed of approximately 30 µm/s for an average duration of 0.77 s. Tumble events occur for an average duration of 0.11 s with a 69*^◦^* average tumble angle. Both values are similar to that of *E. coli*. When exposed to serine, *SM3* suppresses the frequency of tumble events, which in turn increases the average run duration. In addition, the average tumble angle was found to decrease in response to serine. However, when exposed to up to a millimolar concentration of aspartate, *SM3* does not demonstrate a notable change in run-and-tumble parameters. These results suggest that run-and-tumble is the characteristic swimming behavior of *SM3* in its planktonic state. These findings serve as a benchmark in a quest to determine the connection among swimming, swarming, and the complex dynamics of the gut microbiome.

**IMPORTANCE:** Bacteria form the largest domain of living creatures on this planet. Our interactions with bacteria influence us in many ways, not the least of which being in regards to human health. A recently identified species of gut bacteria, *Enterobacter sp. SM3*, has been shown to reduce intestinal inflammation, suggesting that swarming could play a physiologically beneficial role. In this report, we study the motility of individual *SM3* bacteria This study is an essential step towards an overarching goal to understand the influence of bacterial motility on human health.

## INTRODUCTION

Many bacteria have been observed to swim through their environments by rotating an organized bundle of independent, motor-driven, proteinaceous, helical filaments called flagella [1, 2, 3, 4]. A manifestation of this swimming mode, known as run-and-tumble, is characterized by periods of smooth translation at constant velocity, or runs, punctuated by shorter intervals of erratic cell body rotation with little translation, called tumbles. Specifically, runs occur during periods of counterclockwise (CCW) rotation of all flagella, resulting in a helical bundle that pushes the cell body forward. A tumble occurs when one or more flagella switch to the clockwise (CW) rotation state and leave the bundle [5, 6, 7]. The repeated cycles of runs and tumbles result in trajectories resembling that of a random walk. In the presence of a spatial gradient of chemical or environmental stimuli, the run-and-tumble behavior can appear as a biased random walk as the run length towards favored locations can be prolonged to yield net migration. The macroscopic outcome is known as chemotaxis, a crucial microbial behavior that accompanies the run-and-tumble swimming motility [8, 9, 10, 11].

Run-and-tumble has been identified in many peritrichous bacterial species [12, 13, 14, 15]. Among swimming bacteria observed for their run-and-tumble and chemotactic characteristics, *Escherichia coli* has been the most extensively studied with tracking experiments. In the absence of stimuli, the *E. coli* wild type AW405 runs for 1.0 s, tumbles for 0.1 s, and changes direction between successive runs by *∼*68*^◦^*, on average [16, 17, 18]. Furthermore, *E. coli* is shown to be highly responsive to serine, suppressing tumbles and increasing run times by over 300% of those of the control [16]. However, upon testing the response of *E. coli* to aspartate, another chemotactic agent, the effect was found to be insignificant.

*Enterobacter sp. SM3* is a swarming bacterium that has been recently isolated from mouse feces and has been found to reduce intestinal inflammation in mice suffering from symptoms of inflammatory bowel disease (IBD) [19]. Furthermore, the swarming capability of *SM3* has been shown to be directly related to its ability to ameliorate symptoms associated with IBD [19]. The relationship between swimming and swarming is a subject of ongoing studies [12, 20, 21, 22, 23, 24]. However, a first step towards understanding the swarming behavior of *SM3* has to begin with measuring their motion as individual swimmers. In this work, we determine the key run-and-tumble parameters of *SM3* such as the run time, swimming speed, tumble time, and tumble angle. These parameters are measured in a control environment as well as upon exposure to bulk serine or aspartate, which allow for useful comparisons with *E. coli*. We report the occurrence of chemotactic bias in response to serine, including suppression of tumble events, extended run times, and decreased tumble angles as a function of increasing serine concentration, but little bias in response to aspartate. We thus find that *SM3* exhibits swimming behavior notably similar to that of *E. coli*. These initial characterizations can prove useful in further studies involving the swarming behavior of *SM3* and its effect on the intestinal microbiome.

## RESULTS

### *SM3* Exhibits Run-and-Tumble Behavior

The swimming motility of *SM3* was observed using phase-contrast microscopy using a 40x objective in brightfield. Under phase contrast, cells run in relatively straight segments and occasionally change direction with a brief, erratic motion. To better understand the kinematics of these direction-changing events, we examine images taken at a frame rate of 100 FPS. The top half of Figure 1 shows one such representative event as an overlay of images taken seven frames apart. It was observed in real-time that the cell moved horizontally, from left to right, with its body axis parallel with the direction of the motion. Then, during a sequence of 12 frames corresponding to a time of 0.12 s (refer to the bottom half of Figure 1), the cell ceased to translate and rotated by *∼*63*^◦^*. Following rotation, the cell then resumed its motion along a new trajectory *∼*63*^◦^* with the horizontal with its body axis parallel with the direction of motion. The cell shown in Figure 1 rotated continuously in the counter-clockwise direction. However, it was observed that cells can alternate rotation direction during the moment of little translational motion. Through repeated observations, it was found that direction-changing events in *SM3* are characterized by a significant decrease in translation as well as notable rotations of the cell body. These observations are consistent with run-and-tumble behavior; with periods of directed cell motion being called runs and direction-changing events, tumbles.

**FIG 1.**
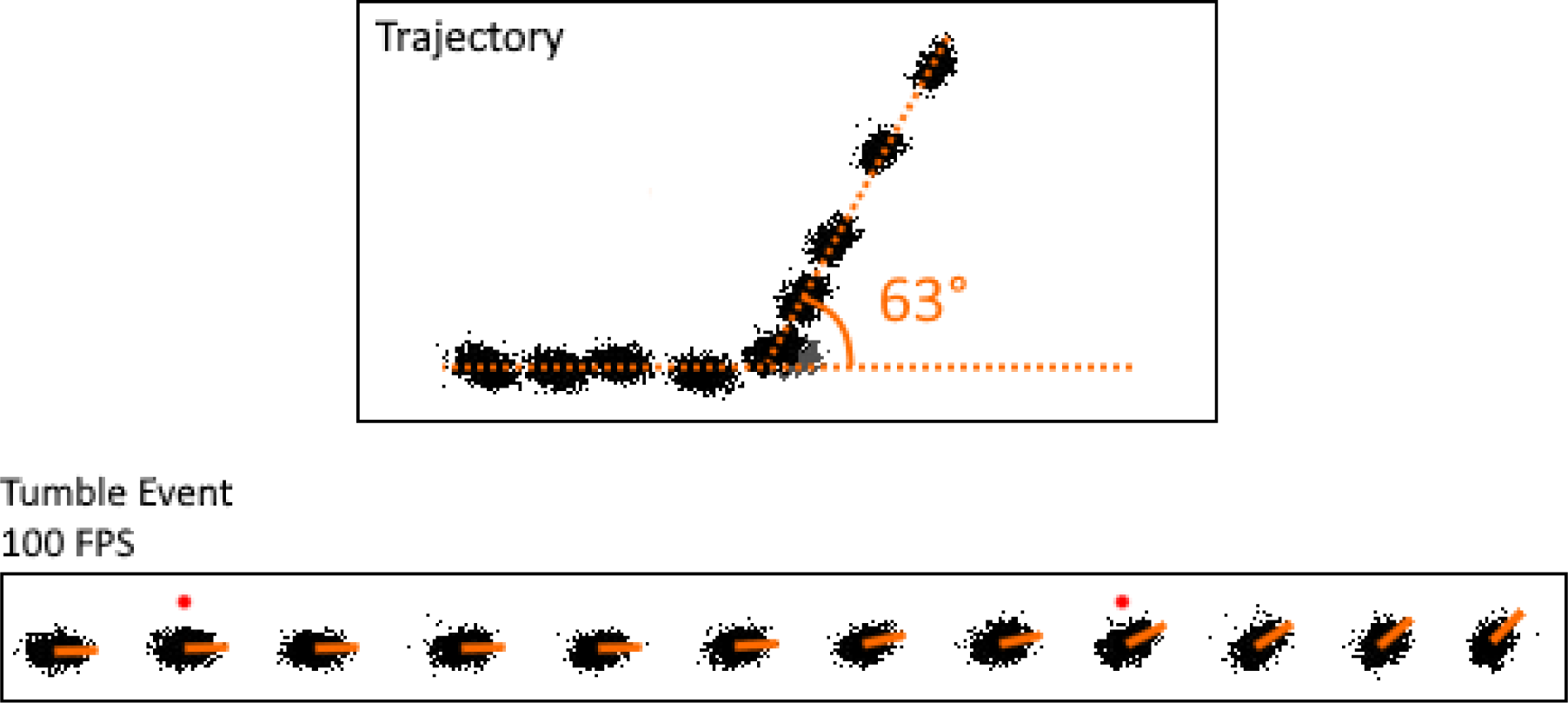
(*Top*) A segment of a *SM3* trajectory. The segment was taken at 100 frames per second (FPS) for 0.7 s resulting in 70 consecutive frames. Images of the cell shown were selected seven frames apart for visual clarity. (*Bottom*) Frame-by-frame sequence of the tumble event from the above trajectory. The event lasts 12 frames, corresponding to a tumble time of 0.12 s. The tumble angle is 63°. Red dots denote two frames that are included in the overlaid image above; they overlap at the position of the tumble. A short movie from which this particular tumble event was captured is provided in the supplemental materials (Movie S1).

### The Run-and-Tumble Motility of *SM3* Is Similar to *E. coli*

To get a quantitative measure of the run-and-tumble motility of *SM3* we employed a custom tracking algorithm. *SM3* was made to swim in a psuedo-2D environment of *∼* 20 µm thick simple motility buffer and movies were acquired using a 10x objective in darkfield. One movie could yield multiple continuous trajectories for analysis. Using the algorithm, a total of 209 tumbles and 199 runs were obtained from 10 separate trajectories. Figure 2*A* shows a representative trajectory of a single *SM3* cell. The trajectory is broken up into relatively straight segments of runs flanked by adjacent tumbling events. The algorithm detected tumble events in the raw position data as instances of decreased linear speed occurring in the neighborhood of increased angular speed. Figures 2*B* and 2*C* plot the linear and angular speed, respectively, as functions of time for the cell whose trajectory is shown. Figure 2*D* shows the tumble events detected by the algorithm denoted as blue lines marking the coincidence of dips in linear speed with peaks in angular speed. These tumble events are also labeled on the trajectory in Figure 2*A* with red dots. The algorithm was applied in a similar fashion to all other trajectories according to the procedure outlined in the Materials and Methods.

**FIG 2.**
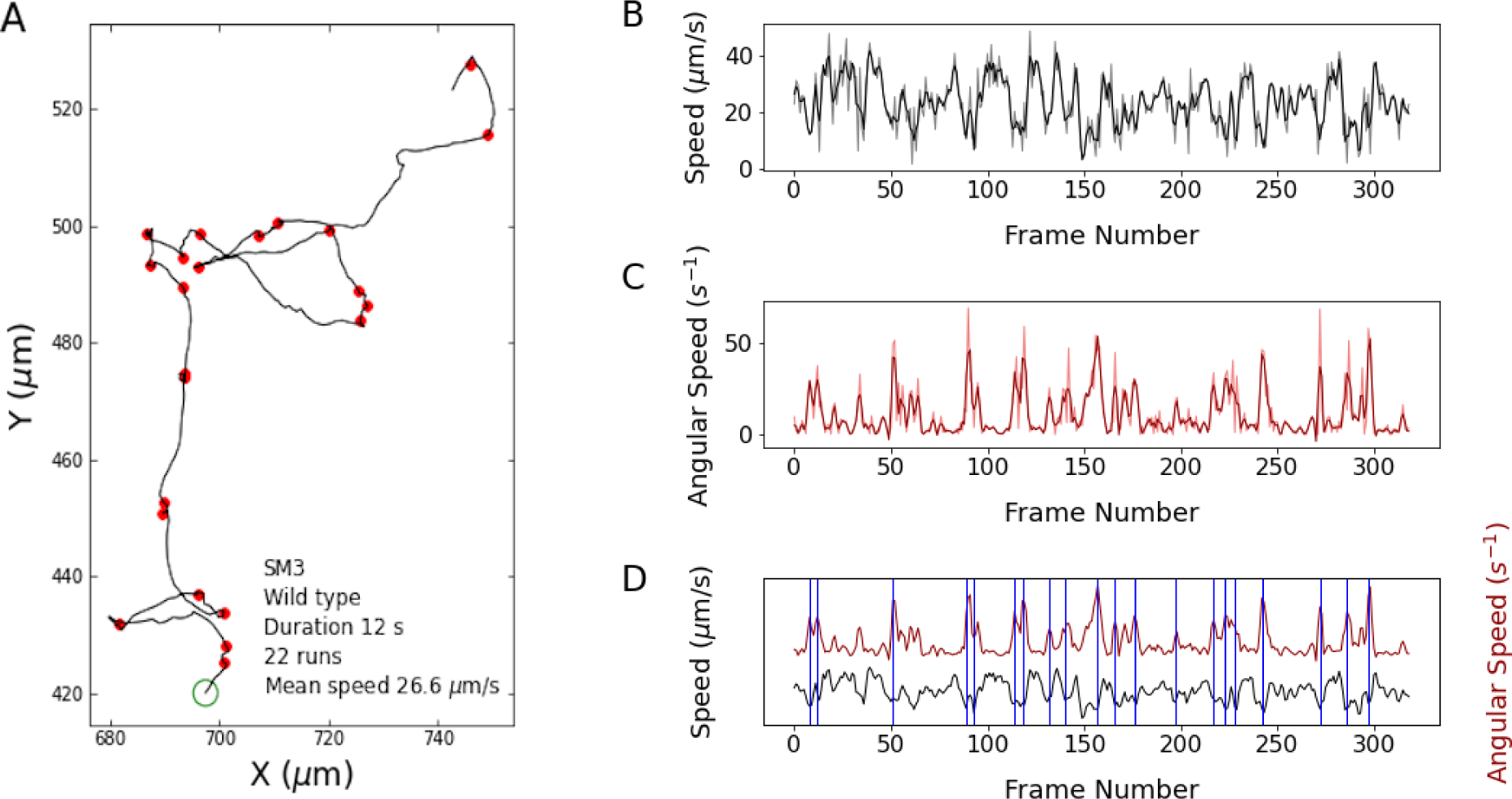
Analysis of *Enterobacter sp. SM3* trajectories performed using a tracking algorithm. (*A*) A representative trajectory. The green circle marks the beginning of the trajectory. The red dots represent tumble events detected by the algorithm. An animated movie of this trajectory is provided in the supplemental materials (Movie S2). (B) Linear speed versus frame number. (C) Angular speed versus frame number. Opaque curves in *B* and *C* show the unsmoothed data, with the solid lines after smoothing. (D) Overlay of the linear and angular speeds on the same plot. Blue lines note the alignment of a dip in the smoothed linear speed with a peak in the smoothed angular speed, indicating a tumble event.

Analysis of several dozen trajectories yield histograms of the obtained tumble times, run times, and tumble angles (Figure 3). The histogram of tumble times follows a bell-shaped distribution with a longer tail on the right and peaked around an average of 0.11 ± 0.05 s. The histogram of run times follows an exponential distribution that drops off for higher values of run time. Interestingly, there is a large percentage of smaller runs with 52% of runs being less than or equal to 0.44 s. The average run time is 0.77 s ± 0.87 s. Additionally, *SM3* runs with an average speed of approximately 30 µm/s and tumbles with an average angle of 69°.

**FIG 3.**
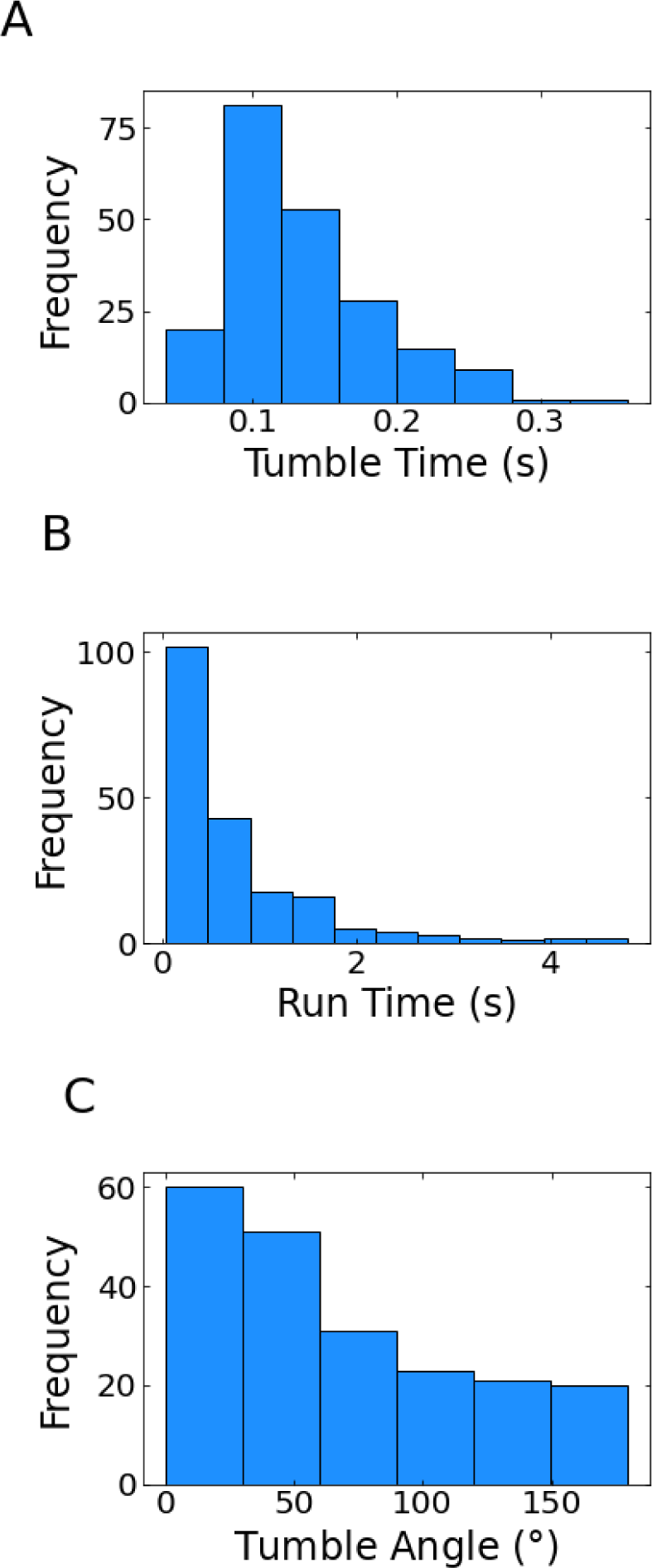
Histograms of the measured (*A*) tumble times, (*B*) run times, and (*C*) tumble angles. The bin numbers selected for the histograms are 9, 11, and 6, respectively.

### *SM3* Alters Its Run-and-Tumble Behavior in Response to Chemoattractants

We exposed *SM3* to two different amino acids, serine and aspartate, and measured its response to them as chemoattractants using the same tracking algorithm. *SM3* shows a strong response to serine, as shown in Figure 4 with data listed in Table 1. The average run time increases with increased serine concentration whereas the average tumble angle decreases. The tumble time (Figure 4*A*) and average speed (Figure 4*B*), however, remain mostly constant over the range of serine concentrations.

**FIG 4.**
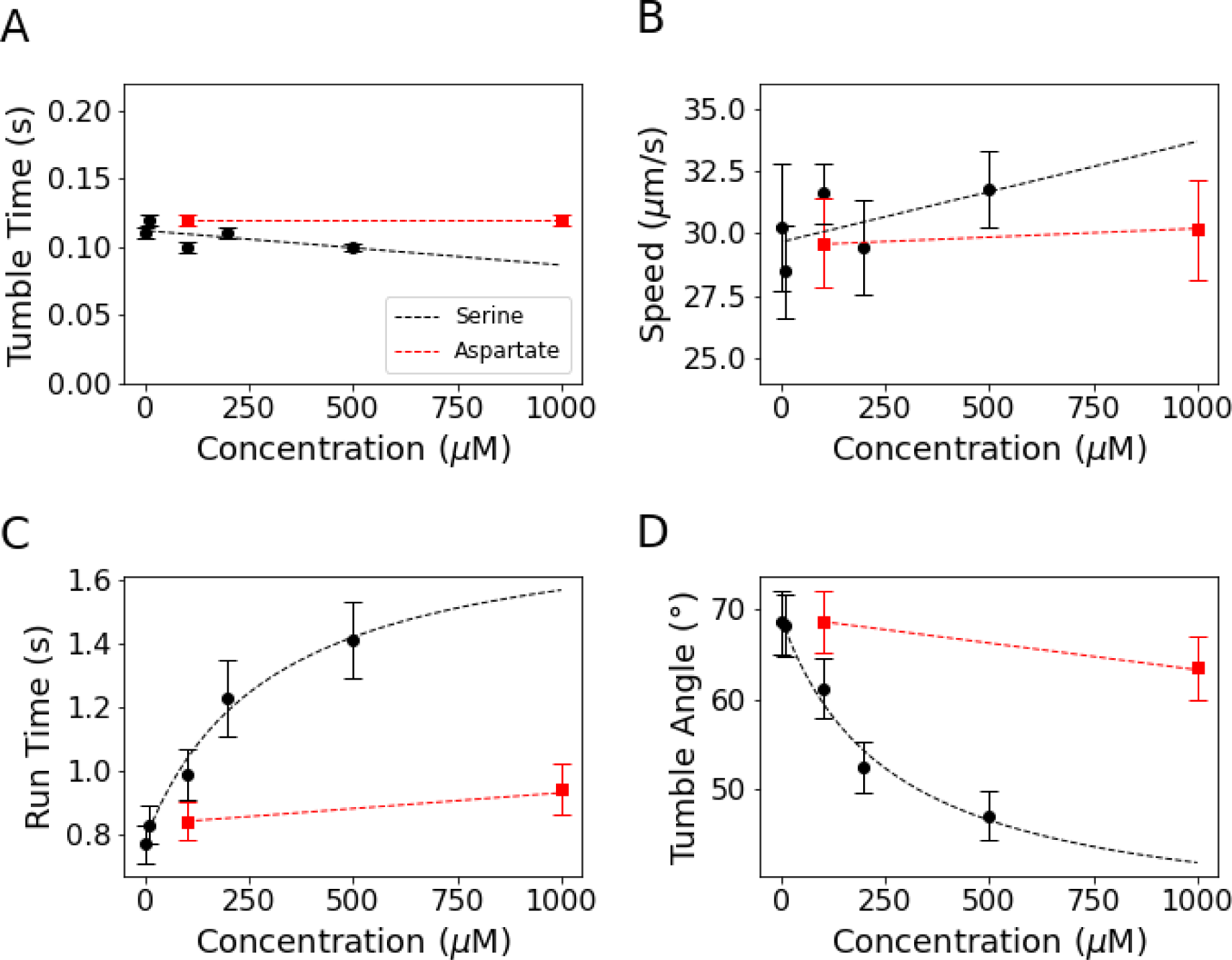
The effect of serine and aspartate on the run-and-tumble behavior of *SM3*. Plots include (*A*) average tumble time, (*B*) average run time, (*C*) average tumble angle, and (*D*) average cell speed as functions of concentration. Dotted lines serve to guide the eye to apparent trends. Error bars are standard errors to the mean.

**TABLE 1.**
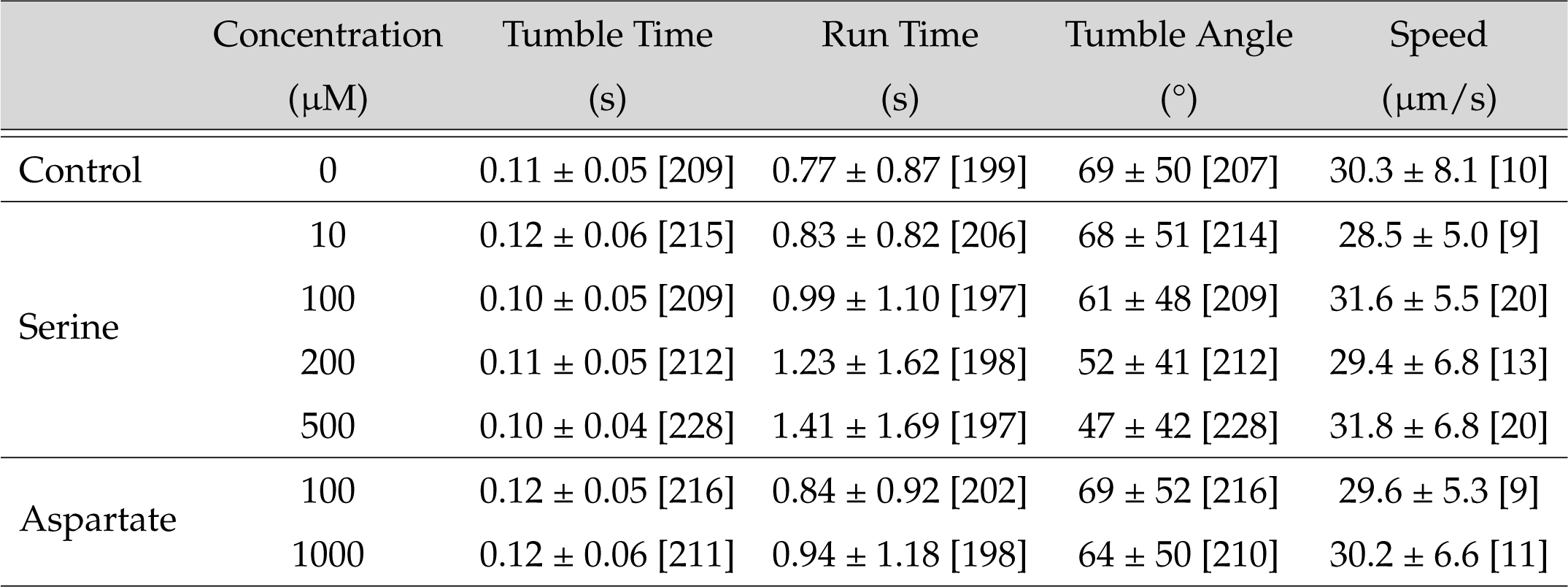
Run-and-tumble analysis of *SM3* swimming in the control environment and upon exposure to serine and aspartate. Tabulated data includes the average tumble time, run time, tumble angle, and speed based on the number of data points shown in brackets. Reported errors are standard deviations.

To test if *SM3* had a preference for the type of chemoattractant in the environment, we exposed it to aspartate in comparison with serine. We chose to use concentrations of 100 µM and 1 mM aspartate, which for serine led to large changes. For both concentrations, aspartate was found to have an insignificant effect on run-and-tumble parameters when compared to the response of serine (Figure 4 and Table 1). These results indicate that aspartate has negligible effects on the run-and-tumble behavior of *SM3* when compared to that of serine. In short, *SM3* exhibits preference for specific chemoattractants.

## DISCUSSION

The data acquired in this study show that *SM3* demonstrates robust motility in its planktonic state with features closely resembling that of the run-and-tumble behavior initially discovered for *E. coli* [16, 18]. *SM3* swims in trajectories that consist of straight runs separated by short tumbles during which the cell body slows down and reorients itself in the direction of a new run (refer to Figures 1 and 2). The average tumble time of *SM3* in LB medium of 0.11 seconds is very similar to that of *E. coli* wildtype AW405, which has an average tumble time of approximately 0.1 seconds. The average tumble angle of *SM3* of 69*^◦^* is also consistent with *E. coli*, which has an average tumble angle of 68°. The average run interval of *SM3* of 0.77 s is only slightly shorter than that reported for *E. coli*, that being approximately 1.0 s. The histogram distributions of tumble time and run time also closely resemble those of *E. coli* with the tumble time distribution being bell-shaped and the run time distribution being exponential [18]. We thus conclude that *SM3* has similar run-and-tumble parameters to that of *E. coli* in simple motility medium.

The response of *SM3* to serine is also similar to *E. coli*. It has been demonstrated in the literature that serine suppresses the frequency of tumble events in *E. coli*, resulting in the lengthening of its runs. *E. coli* also shows no response to aspartate [16, 18]. An interesting finding, though, is the tendency of *SM3* to decrease its average tumble angle as the bulk serine concentration increases (refer to Figure 4). This decrease in the average tumble angle gives rise to smoother trajectories of *SM3* in the presence of a chemoattractant. This inverse relationship of tumble angle to the concentration of chemoattractants, to our knowledge, has not been reported in the literature. Biologically, however, this behavior makes sense: when bacteria detect that the local concentration of chemoattractants such as serine have increased, it may be preferable to continue moving in the direction of travel even if interrupted by a tumble.

An explanation for the angle dependency of *SM3* on serine requires a microscopic understanding of tumble kinematics. Tumble events were observed to be characterized by reduced swimming speed and noticeable reorientation of the cell body (see Figure 1). A tumble is initiated when at least one flagellum switches from the CCW to the CW state [6]. We hypothesize, however, that most often a single switch is responsible for a tumble event. The average run time is on the order of 1 second for both *E. coli* and *SM3*. If every motor can switch with equal probability and if one switch suffices to end a run, the rate of a CCW-CW switch must be lower than on the order of 1 per second. If one further assumes that the motors on the cell body are not correlated with one another in their switches [25, 26, 27], the probability of two or more motors switching to the CW state during the short interval of time associated with a tumble (lasting on the order of 0.1 sec) is lower by a factor of ten, approximately. Due to the high rotation rate of the motor (on the order of a hundred turns per second), however, abruptly switching the rotation direction of a single flagellum may cause more than one flagellum to leave the bundle of several flagella, which were hydrodynamically coupled during the last run (refer to Figure 5). Other flagella may also immediately splay apart and become uncoordinated, causing the cell to cease swimming and start to tumble. During the tumble, cell body reorientation is driven by the net torque on the cell body due to forces exerted by both the CCW and CW rotating flagella. The time interval of the tumble is brief enough such that the net torque on the cell body may not change much in direction. Therefore, cell body reorientation is expected to be continuous, as captured by the 100 FPS images shown in Figure 1. When the stray flagellum switches back to the CCW state, the tumble ends and a new run begins as the bundle reforms. This model of tumble kinematics, although explaining the behavior we observed for *SM3*, requires further experimental confirmation. Studies aided by fluorescent imaging of live flagella are necessary to clarify the kinematics and the actual physical mechanism of the tumble event for *SM3*, as previously done for other peritrichous bacteria such as *E. coli* [6, 28].

**FIG 5.**
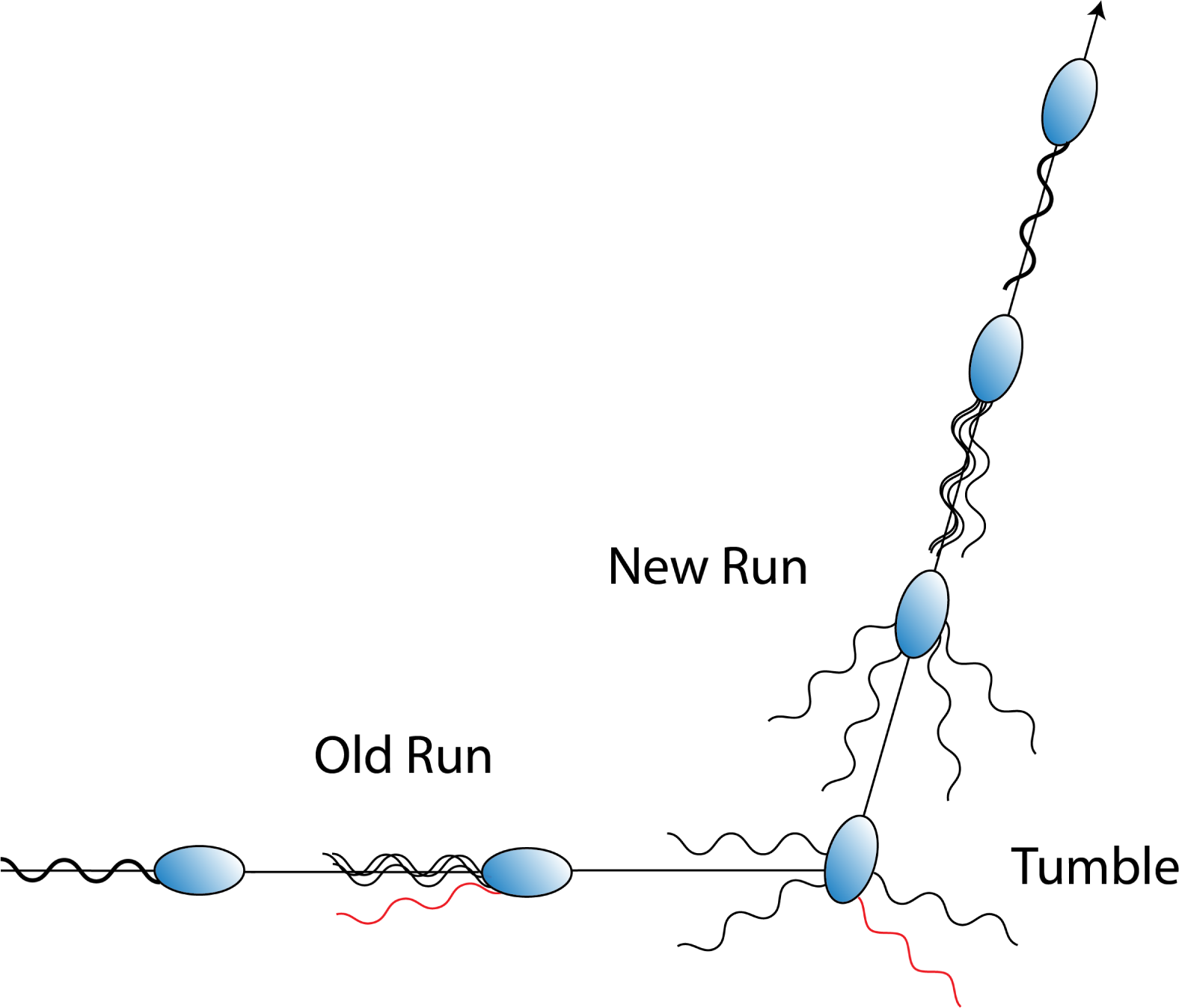
Diagrammatic representation of a tumble event in *SM3*. When one flagellum switches rotation direction from CCW to CW, it leaves the bundle and causes the bundle to become uncoordinated, initiating a tumble. During the tumble, motility is reduced and the cell reorients itself. This diagram can be compared with the observed behavior in Figure 2. Flagella were not observed directly in this study.

A possible mechanism for the inverse relationship between the average tumble angle and the concentration of serine is the reduction of the average tumble time. Although the measured data show that the average tumble time only decreases slightly over a large range of serine concentrations (refer to Figure 4*A*), the results may be limited by the time resolution of the data acquired. Since most of the movies were taken at 25 FPS, tumble times determined by the algorithm are multiples of 0.04 s. For example, if one hundredth of a second difference in the average tumble time could cause a significant change in the average tumble angle, that difference, as indicated by the trend shown in Figure 4*A*, would not be reliable given the margin of error of our measurements. Other possible mechanisms would require a more detailed investigation of the mechanical properties of the flagellum motor as well as direct imaging of the flagella themselves, which goes beyond the scope of this report on the first study of *SM3* swimming motility. Future investigations are necessary to explore the run-and-tumble behavior further for *SM3*, particularly in its swarming state.

In conclusion, we report an analysis of the swimming behavior of the newly identified species of gut bacteria *Enterobactor sp. SM3*. We find that *SM3* exhibits a swimming motility that resembles the run-and-tumble behavior of *E. coli*. Using an algorithm that associates tumble events with periods of low translational speed and high angular speed, we reliably extracted key run-and-tumble parameters. In response to increasing amounts of bulk serine and aspartate, both the average tumble time and cell speed remained constant. However, in the presence of serine, the average run time increased while the average tumble angle decreased. We proposed a plausible mechanism, which requires confirmation with data of better time resolution, as well as further studies involving measurements of motor rotation frequency or direct imaging of live flagella as the concentration of serine is varied.

## MATERIALS AND METHODS

### Preparation of Bacterial Cultures

*Enterobacter sp. SM3* cells were extracted from a −80 °C frozen stock using a sterilized inoculation loop and inoculated in a 10 mL solution of lysogeny broth (LB) medium (10 g/L tryptone, 5 g/L yeast extract, and 5 g/L NaCl dissolved in de-ionized water). Cells grew overnight at 37 °C in an incubator shaking at 160 rpm. From the overnight growth, 100 µL of bacterial culture was transferred using a micropipette to 10 mL LB for a fresh growth. This was placed in the shaking incubator to resume growth for four hours, which yielded optimal swimming motility for optical microscopy [29].

Prior to microscopy, 10 µL of fresh growth was transferred to a 1 mL Eppendorf tube and mixed with 990 µL of a simple motility buffer containing 67 mM NaCl, 0.1 mM EDTA, and 10 mM sodium phosphate at pH 7.0.

For chemotaxis studies, a 0.1 M stock solution of either l-serine or l-aspartic acid monosodium salt in de-ionized water was made and all subsequent concentrations of each chemoattractant were made by diluting a stock solution into the motility buffer. 10 µL of a chemoattractant solution at varying concentrations was added to 1 mL of control solution consisting of 990 µL of buffer and 10 µL of bacterial solution.

### Slide Preparation, Imaging, and Processing

The perimeter of a glass coverslip (22 mm x 22 mm) was coated with a thin layer of vacuum grease. In the center of the coverslip, 5 µL of bacterial solution was deposited and a glass microscope slide (3" x 1" x 1 mm) was gently pressed on top until the fluid formed a thin, uniform layer with an estimated thickness in the range of 10-20 µm. The slide was then transferred to the stage of an inverted optical microscope with the coverslip facing down.

A Nikon Eclipse TE2000-U inverted optical microscope was used for microscopy measurements. Movies were taken using a Point Grey GS3-U3-51S5M-C Grasshopper3 CCD camera with a resolution of 2448 x 2048 and a pixel conversion factor of 1 pixel = 0.342 µm, which was determined using a calibration slide. The software Point Grey Fly Capture 2 was used to record frames in .tiff format. For experiments requiring high spatial and temporal resolution to observe tumble events, a 40x Nikon objective was used to capture brightfield images at 100 FPS. For run-and-tumble experiments using the algorithm, a 10x objective was used to capture darkfield images at 25 FPS.

Frame-by-frame images were processed using the open-source software, Fiji. The images were loaded into Fiji and the software was calibrated using the pixel conversion factor listed above. The images were then converted to 8-bit format and thresholded using a maximum entropy filter; using the dark background setting for darkfield images. In this way, nicely contrasted images with bacteria appearing black on a white background were produced, suitable for effective cell tracking.

### Cell Tracking

Cells were automatically tracked using the Trackmate package in Fiji. Trackmate is a free plugin which allows for easy and efficient semi-automatic tracking of bacteria over time. The program implements tracking in two main steps: cell detection and linking [30, 31]. In the cell detection step, the user can choose from a selection of different detectors. For our experiment we chose to implement a difference-of-Guassian (DoG) detector. Prior to spot detection, the DoG detector applies a DoG filter to the 8-bit images to approximate a Laplacian of Guassian (LoG) filter. This serves two purposes: to blurr the image, and therefore reduce high-frequency spatial noise, and to enhance the edge boundaries of cells, making them easier to detect by the program [32]. After filtering, the program detects cells subject to two main criteria: the approximate cell diameter and the quality threshold. For *SM3*, the average cell length was found to be 1.5 *±* 0.3 µm. This was determined by measuring the long axis of 121 separate bacteria using the length tool in Fiji and taking the average. Thus, we were able to supply the program with a reasonable size around which an object could be identified as a bacterium. The other parameter that the user can modify is called the quality threshold and is a measure of the quality of the spots detected. Spots below the designated quality threshold are filtered out by the program to reduce the detection of spurious spots. The threshold is determined by trial and error. Once the algorithm has been optimized and cells are detected, cell linking can begin.

Trackmate provides multiple trackers to the user depending on the system of interest. For our experiment, we used a simple linear assignment problem (LAP) tracker to link our detected cells into a cohesive track. The simple LAP tracker works by assigning a cost to linking two detected cells at time t and t + Δt (i.e. between consecutive frames) which is proportional to the squared distance between the objects. The tracker assigns links in such a way as to minimize the total linking cost [33, 34]. Simply put, the tracker does its best to link cells between frames that are close to one another in position. There are, however, factors that can disrupt the linking process [31]. For instance, cells can move in and out of the microscope’s plane of focus, causing gaps, and they can collide with other cells or anomalous spots, causing confusion for the tracker. To circumvent these issues, we adjusted three key parameters. The first parameter, called the max linking distance, is the minimum distance over which the tracker looks for candidate cells to link. If the max linking distance is set to X µm, then cells in frame t + Δt must be within X µm of the cell in frame t to be considered as part of that cell’s trajectory. For our cells, with an average speed of 30 µm/s and a time interval between frames of 0.04 s, a max linking distance of 1-2 µm would be reasonable. We chose to give the tracker more freedom and set the max linking distance to 5 µm. The second parameter of interest, the gap-closing max distance, is the maximum distance the user allows the tracker to bridge a gap between two track segments. This preserves the track in the event of a cell disappearing from the plane of focus for multiple frames. We set this parameter again to 5 µm, or a little over three cell body lengths. It should be noted, however, that in selecting bacteria to track we went out of our way to avoid those trajectories which exhibited noticeable out-of-plane motion. The last parameter that we adjusted is called the gap-closing max frame gap. It is the maximum frame interval between which two track segments will be bridged. We chose this value to be two frame intervals or 0.08 s. With these parameters for detection and linking, we were able to effectively produce long, reliable trajectories of the in-plane motion of *SM3* as shown in Figure 2*A*. All data were exported from Fiji in .csv format to be analyzed by our custom algorithm. The exported data included the (X, Y) positions and frame numbers of the tracked cells.

### Analysis of Cell Trajectories

Trajectories were chosen not to include tethered cells, dead cells, and other external interactions that could influence bacterial movement. Cell trajectories were analyzed using a custom event recognition algorithm (github.com/Tang-BioPhys/SM3-Motility) based on the work of Masson *et al.* [35] and repurposed by *Theves et al.* [36]. This algorithm provides a simple and concise means for extracting several key run-and-tumble parameters such as tumble time, run time, tumble angle, and cell speed. We used the kinematic equations

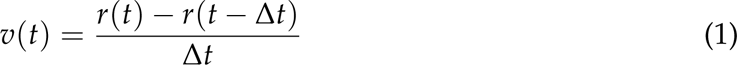

and

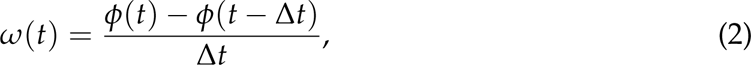

where *ϕ* is determined by representing the linear velocity in polar coordinates, e.g.,

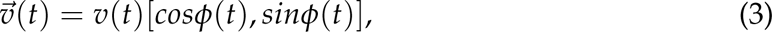

to calculate the semi-instantaneous linear and angular speeds as functions of time. For our experiment, Δt = 0.04 s, given the frame rate of 25 FPS.

The linear and angular speed raw data were passed through a Savitsky-Golay filter for smoothing. It was found empirically that a window size of five and a polynomial order of two yielded the best smoothing while minimizing the amount of lost information. Refer to Figure 2*B* and 2*C* in the Results section for representative plots of the linear and angular speed as functions of time, both smoothed and unsmoothed.

The algorithm automatically detected local minima in the smoothed linear speed data and, of those minima, selected for tumble events. If the depth satisfied the requirement,

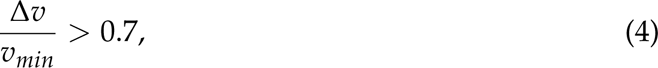

where Δv is the depth and v_min_ is the minimum speed, then the minimum was recorded by the program as a possible tumble event. The value of 0.7 is empirical and was found to detect tumbles consistent with those noted by the naked eyes. Once all potential tumbles were detected, the algorithm calculated the tumble time corresponding to each minimum by counting the frames of the neighboring speeds around v_min_ as part of the tumble if and only if:

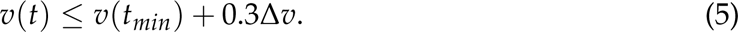

Again, the choice of threshold was empirical and we found that 0.3 returned results most consistent with observation.

In addition to the criterion given by Equation 4, a sufficient increase in the cell’s angular speed, in the neighborhood of a decrease in linear speed, was required for an event to be considered a tumble. Figure 2*D* in the Results shows a superimposed plot of the linear and angular speed as functions of time. If a dip in the linear speed corresponded with a peak in the angular speed that was greater than 20% above the baseline, then the event was considered a verified tumble. These events are indicated by the blue lines in Figure 2*D*.

The advantage of this approach is two-fold. First, it prevents false-positives by requiring that both speed criteria be met. However, because of the requirement that there be a sufficient decrease in the cell speed, the algorithm is prone to missing certain tumble events. For example, the algorithm may not detect events in which the cell rapidly changes its angle but does not show a significant change in speed. This manifests itself primarily when the timescale of the tumble event is less than the frame interval of our camera (0.04 s). However, we found such events to be rare. By requiring that both the linear and angular speed criteria be met, we ensure that all detected tumble events are valid at the expense of missing short tumbles on rare occasion. In addition, this approach filters out anomalies that may have passed the criterion given by Equation 4. For instance, events in which the cell slows down or even stops without showing a significant change in angular speed. These events should not be confused with the form of bacterial motility known as pausing which has been reported for other species of bacteria [37, 38, 39]. In short, we found that the algorithm effectively and reliably identifies tumble events.

Runs are defined as long periods of uninterrupted motion in which a cell moves at roughly constant speed along its long axis between two tumble events. The algorithm determines the run times by summing the intervals of frames between adjacent tumbles. Tumbles that do not meet both criteria above are considered to be part of a larger run. Once the tumble and run times have been determined, other relevant parameters are readily determined by the algorithm. The average cell speed is calculated by identifying long runs in a given trajectory (defined as any run lasting six frames or more), assigning an average speed to each run segment, and then averaging those values to assign an average speed to a given trajectory. Therefore, the number of speed data points equals the number of analyzed trajectories. The average tumble angle was determined by using the average incident and outgoing angle of the trajectories. The incident and outgoing trajectories were taken over three frames before and after the tumble event, respectively. Since the tumble angle is defined by a directional change in the trajectory, and not the cell body orientation, it is unaffected by rotations of the cell body into the third dimension during a tumble, as long as the trajectory itself remains pseudo-2D. As a result, we obtained a large data set of tumble times, run times, tumble angles, and cell speeds under each solution condition.

## ACKNOWLEDGMENTS

Dr. Sridhar Mani at the Albert Einstein College of Medicine kindly provided the bacterial strain, *Enterobacter sp. SM3*, and offered comments to the manuscript. We thank Dr. Thomas Powers at Brown University for valuable discussions.

This work was funded by the National Science Foundation (NSF DMR-2207284).

